# MetageNN: a memory-efficient neural network taxonomic classifier robust to sequencing errors and missing genomes

**DOI:** 10.1101/2023.12.01.569515

**Authors:** Rafael Peres da Silva, Chayaporn Suphavilai, Niranjan Nagarajan

## Abstract

**Background:** With the rapid increase in throughput of long-read sequencing technologies, recent studies have explored their potential for taxonomic classification by using alignment-based approaches to reduce the impact of higher sequencing error rates. While alignment-based methods are generally slower, k-mer-based taxonomic classifiers can overcome this limitation, potentially at the expense of lower sensitivity for strains and species that are not in the database.

**Results:** We present MetageNN, a memory-efficient long-read taxonomic classifier that is robust to sequencing errors and missing genomes. MetageNN is a neural network model that uses short k-mer profiles of sequences to reduce the impact of distribution shifts on error-prone long reads. Benchmarking MetageNN against other machine learning approaches for taxonomic classification (GeNet) showed substantial improvements with long-read data (20% improvement in F1 score). By utilizing nanopore sequencing data, MetageNN exhibits improved sensitivity in situations where the reference database is incomplete. It surpasses the alignment-based MetaMaps and MEGAN-LR, as well as the k-mer-based Kraken2 tools, with improvements of 100%, 36%, and 23% respectively at the read-level analysis. Notably, at the community level, MetageNN consistently demonstrated higher sensitivities than the previously mentioned tools. Furthermore, MetageNN requires *<* 1*/*4^*th*^ of the database storage used by Kraken2, MEGAN-LR and MMseqs2 and is *>*7x faster than MetaMaps and GeNet and *>*2x faster than MEGAN-LR and MMseqs2.

**Conclusion:** This proof of concept work demonstrates the utility of machine-learning-based methods for taxonomic classification using long reads. MetageNN can be used on sequences not classified by conventional methods and offers an alternative approach for memory-efficient classifiers that can be optimized further.

## 1 Background

In recent years, there have been rapid advances in our understanding of microbial diversity, the communities they form, and their impact on human health [1]. These have been enabled by the widespread use of high-throughput sequencing technologies, particularly by leveraging *metagenomics* to directly analyze a diverse pool of DNA and circumvent the limitations of microbial culture [2]. The resulting data then needs to be computationally deconvolved to study the genetics of the organisms that gave rise to the DNA pool, with many bioinformatics tools designed to balance sensitivity, precision and resource-intensiveness of the analysis [3].

Metagenomic analysis workflows follow one of two different paradigms: (i) *de novo* assembly and binning tools that help uncover genomes for further investigations when no prior knowledge of a microbial community is available, versus (ii) taxonomic classification tools that match reads against a reference database and are used in applications where suitable references are available (e.g., pathogen detection), or where sequencing and analysis costs need to be reduced (e.g., for complex communities or large-scale clinical studies) [3, 4, 5]. The use of these paradigms is also dictated to an extent by the sequencing technologies employed, with taxonomic classification being more popular for short reads (*<*300bp, e.g., with Illumina sequencing) while longer reads (*>*1kbp using systems developed by Pacific Biosciences and Oxford Nanopore Technologies, ONT) are favored for *de novo* assembly [6]. As long-read technologies become more cost-effective and widely accessible, recent studies have also explored their promise for highly accurate and sensitive taxonomic classification (e.g., MEGAN-LR [7] and MetaMaps [8]). These have taken an alignment-based approach to reduce the impact of higher sequencing error rates (typically *>*1%) [6, 9]. However, alignment-based tools are generally slow and k-mer-matching tools, such as Kraken2 [10], can be used to overcome this limitation, but at the expense of relying on large databases. Additionally, benchmarking studies suggest that these tools do not generalize well for reads from genomes that are not in the database used, usually returning false positives from distant lineages [4].

An alternative paradigm for taxonomic classification, leveraging the strength of machine learning to generalize across data, is the use of deep-learning in methods such as GeNet [11] and DeepMicrobes [12] which were designed for accurate short reads. While these were innovative, they could not outperform exact k-mer matching tools [11, 13, 14, 15, 12]. The feasibility of designing machine learning-based methods that work with erroneous long reads and generalize well across genomes relative to other taxonomic classification approaches thus remains an open question.

In this work, we present MetageNN, a neural network taxonomic classification method robust to sequencing errors and missing reference genomes. MetageNN overcomes the limitation of not having long-read sequencing-based training data for all organisms by making predictions based on k-mer profiles of sequences collected from a large genome database. We use short k-mer-profiles that are known to be less affected by sequencing errors to reduce the “distribution shift” between genome sequences and noisy long reads. Comparisons using synthetic long reads demonstrate MetageNN’s efficiency and robustness to sequencing errors relative to other deep-learning-based methods for taxonomic classification (GeNet and DeepMicrobes). Furthermore, in benchmarking experiments with ONT reads from bacterial isolates and pseudo-mock communities, MetageNN outperforms alignment (MetaMaps and MEGAN-LR) and k-mer-based (Kraken2) classifiers in detecting potentially novel lineages (i.e. reads from species out of the database). Additionally, MetageNN is *>*7x faster than MetaMaps and GeNet and needs *<* 1*/*4^*th*^ of the memory required by Kraken2, MEGAN-LR and MMseqs2 for database storage.

These results demonstrate the feasibility of machine learning methods for taxonomic classification using long reads, enabling predictions for a large number of taxa with limited sequencing data and providing a sensitive classification of novel lineages while being memory-efficient.

## 2 Results

### 2.1 Setting parameters for the MetageNN model

We conducted several preliminary experiments to establish parameter choices for MetageNN, using the “small database” (described in section Databases) to enable extensive testing.

#### 2.1.1 K-mer size analysis

Considering computational constraints and the fact that short-k-mers are more robust to sequencing errors, we tested k-mer sizes ranging from 3 to 6. Three training datasets were constructed consisting of 2000, 4000 and 6000 sequences per genome (1kbp sequences) and 200 sequences per genome were sampled for the validation dataset. MetageNN was trained on each dataset until convergence was reached on the validation dataset (early stopping to prevent overfitting). In order to assess k-mer robustness to sequencing errors potentially found in long-read settings, we introduced noise profiles similar to ONT sequencing into the validation dataset with the BadReads tool [16], resulting in a synthetic long-read test dataset (the test dataset has an average accuracy of 95% when aligned to reference genomes). As shown in fig. 1 (a), MetageNN results improved with larger k-mers for both error-free sequences and synthetic ONT reads and benefitted from having more sequences for training, and correspondingly we set k-mer size to 6 and further explored the impact of genome coverage.

**Figure 1.**
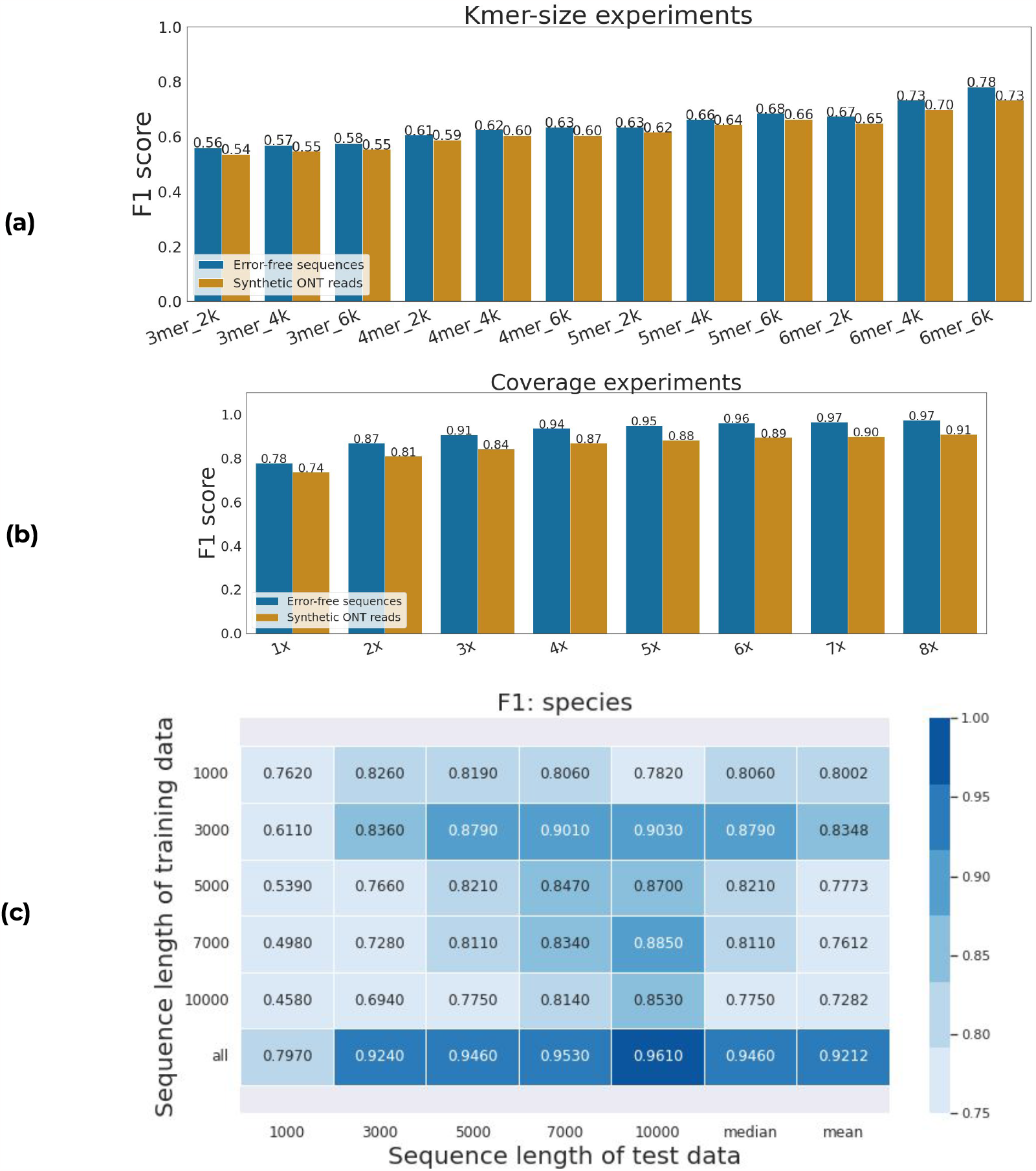
k-mer size, coverage and sequence length analysis. F1 scores for error-free sequences and synthetic ONT reads. (a) MetageNN results by increasing k-mer size and sequence samples. (b) MetageNN results by coverage used. (c) MetageNN results for F1 scores when trained on different sequence lengths. The rows indicate which sequence length the model was trained on. The columns represent the test datasets for each sequence length.

#### 2.1.2 Coverage analysis

As training time could be a limiting factor for this problem, we sought to evaluate the amount of read coverage needed per genome for MetageNN’s performance to saturate. We created eight training datasets with increasing coverage (1*×* to 8*×*) per genome. We report F1 scores on the validation dataset of error-free sequences and synthetic ONT reads dataset (same as in K-mer size analysis). As seen in fig. 1 (b) MetageNN’s performance starts to saturate from around 5*×* coverage in both settings. Therefore, MetageNN trained with a 5*×* coverage and above per genome is a suitable setting when moving to the training using the “main database” (Databases).

#### 2.1.3 Sequence length analysis

To evaluate the impact of sequence lengths on model training and performance, we created five training datasets with fixed sequence lengths of 1, 3, 5, 7, and 10kbp, respectively (1*×* coverage), with validation and test datasets with similar length profiles but fewer reads (200 per genome, error-free sequences).

Training and testing MetageNN models independently for each dataset based on sequence length provided the results summarized in fig. 1 (c). Overall, we found that the highest performance was achieved for test datasets with the same or longer sequences than the one used for training. It is interesting to note that the longer the sequence used for training, the lower the observed performance for shorter test sequences (similar to the trends observed in previous work [17]). Therefore, using a shorter sequence for training seems to have an advantage, potentially due to the sparseness of k-mer-profiles for shorter sequences. We noted that the sequence length bias could potentially be eliminated by training MetageNN using data for all sequence lengths available, resulting in the highest mean and median accuracy results across all test datasets (fig. 1, (c) last row). However, a potential drawback of this approach is its requirement for more sequences and longer training time to ensure adequate performance across different sequence lengths.

### 2.2 MetageNN is more effective and robust to sequencing errors than existing deep-learning-based taxonomic classification methods

Having established MetageNN building blocks (see Setting parameters for the MetageNN model), we move on to benchmarking against existing taxonomic classification methods. First, we aimed to evaluate the efficiency and robustness against sequencing errors of MetageNN relative to existing deep-learning-based taxonomic classification methods.

Existing taxonomic classifiers based on deep learning were trained and tested using short-read data (Genet [11] and DeepMicrobes [12]). Nevertheless, we investigated if these methods can work with long-read data, starting with ideal conditions, i.e., error-free sequences to assess the scalability of training time and classification speed. We also assessed robustness to sequencing errors using synthetic long-read data with different error rates. By using these ideal conditions on the “small database”, we were able to avoid posterior training using millions of sequences from a large database of genomes since some methods may present an unfeasible training experience. We report the time taken to complete one training epoch, the classification speed, and F1 scores on noisy synthetic ONT long reads of MetageNN and existing deep-learning-based taxonomic classification.

#### 2.2.1 Baselines

GeNet [11] is a convolutional neural network (CNN) based taxonomic classification model. In the first step, it employs a one-hot encoding strategy for DNA letters in the sequence, followed by sequencing embedding to learn representations of it, as well as a positional embedding approach (e.g., base positions are concatenated to the one-hot encoding before being used as input to an embedding layer) [18]. The result of these two embeddings is the input to the CNN (ResNet architecture [19]). This strategy is limited by its fixed read length *k*. In contrast to short-reads, which usually present a fixed read length, long reads present read lengths ranging from a few hundred bases to thousands of kilobases [20]. Thus, information from sequences longer than *k* is ignored, while information from sequences smaller than *k* is padded in GeNet.

On the other hand, DeepMicrobes [12] is based on recurrent neural networks [21]. Specifically, this model uses a Bidirectional-LSTM [22] followed by a self-attention layer [23] to learn the feature representations, which are then used as inputs to a multi-layer feed-forward neural network. It encodes its input reads by dividing them into fixed-size k-mers, then embedding each k-mer and using it as input to the Bidirectional-LSTM. Consequently, this approach may present slow training and inference for long reads since its forward and backward gradients’ steps depend on the read length that might reach thousands of bases for long-read sequencing technologies.

#### 2.2.2 Experimental setup

For our training data, we randomly sampled 1*×* coverage of sequences with a fixed length of 1kbp from the 47 genomes found in the “small database” (approximately 235,000 sequences in total).

Validation data was created in the same way as the training dataset, but with fewer samples (200 sequences per genome) and sampling using different seeds. GeNet, DeepMicrobes and MetageNN were trained and tested using the same machine (one GPU). We fine-tuned the baselines using the validation dataset following the author’s recommendations for hyper-parameters in GeNet and DeepMicrobes and trained until convergence (early stopping based on validation loss). MetageNN architecture for these experiments is depicted in fig. 5, a, blue text.

In order to assess the model’s robustness to sequencing errors found in long-read settings, we used the BadReads tool [16] to create synthetic ONT long reads by introducing sequencing errors on the 1kbp sequences from the validation dataset. We generated three test datasets representing scenarios with low, moderate and high sequencing error rates (median accuracy 95%, 90% and 80%, respectively).

#### 2.2.3 Results

fig. 2 summarizes the results. In spite of the different architectures and variety of parameters presented by GeNet (approximately 17 million trainable parameters) and DeepMicrobes (69.8 million trainable parameters), we were interested in how long it would take for these methods to complete their training epoch, given that once an architecture is selected the amount of training data can be substantial for a large database of genomes (millions of sequences). With this, GeNet underwent 48 seconds to complete one epoch round (fig. 2, a). DeepMicrobes is based on recurrent neural networks, which means that for each k-mer it will perform forward and backward propagation. In these conditions, with a sequence length of 1kbp, this method required 243 seconds, potentially making it computationally intractable for longer reads from a large database of genomes (fig. 2, a). Furthermore, metagenomic sequencing experiments can contain millions of reads, thus classification speed (i.e., inference time) is an important metric. We measured the total number of sequences processed per second for existing methods using the same machine (no GPUs used). In this case, GeNet is faster than DeepMicrobes, with 143 sequences per second compared to 19 for DeepMicrobes (fig. 2, b). These results corroborate that existing methods, such as DeepMicrobes or possibly methods based on recurrent neural networks, might not be feasible for long reads. In noisy test datasets (fig. 2, c, d and e), DeepMicrobes reported higher F1 scores than GeNet in all three settings, but at the cost of classification speed and epoch run time (fig. 2, a and b).

**Figure 2.**
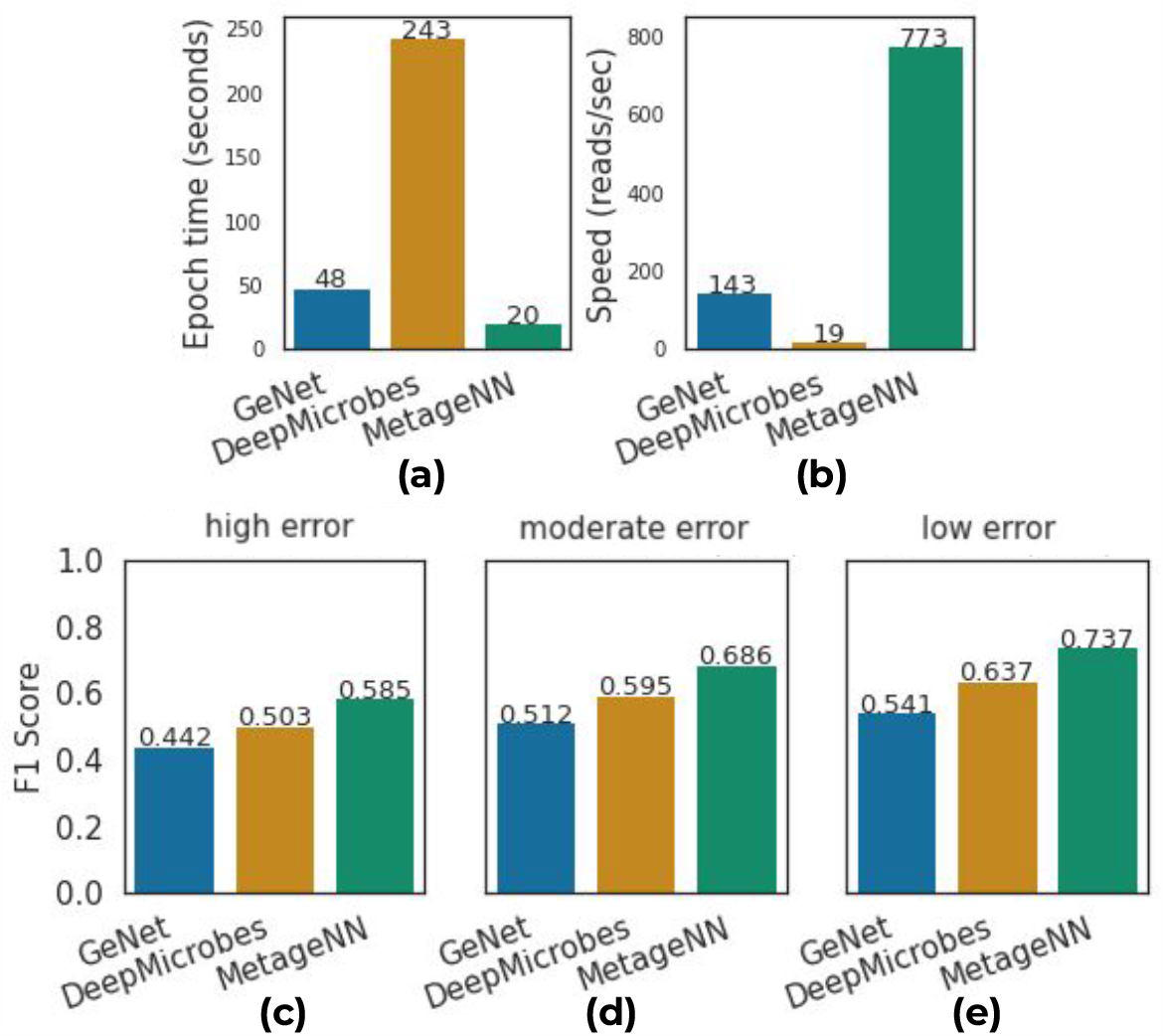
An analysis of existing deep-learning-based taxonomic classification methods and MetageNN for long-read settings. For each method, we (a) show the epoch run time elapsed and (b) display the inference time in reads-per-second. (c)(d) and (e) show the results for F1 score on three test datasets simulating higher, moderate and lower rates, respectively. As compared to existing approaches, MetageNN has the shortest epoch run time, the highest speed and the highest F1 scores demonstrating its robustness to sequencing errors present in long-read data.

Finally, we evaluated MetageNN (approximately 10.7 million parameters) in these same settings. MetageNN is based on short-k-mer profiles obtained from error-free training sequences (fig. 5, a). MetageNN presented the fastest epoch run time of approximately 20 seconds and a classification speed of 773 sequences per second (including k-mer counting time) (fig. 2, a and b). MetageNN also reported the highest F1 scores on the three synthetic ONT test datasets with a varying number of sequencing errors introduced.

### 2.3 MetageNN presents a higher F1 score than GeNet, and it is more sensitive than MetaMaps, Kraken2, GeNet and MEGAN-LR at the read level for bacterial isolates reads of species out of the database

In this section, we proceed to evaluate MetageNN and existing baselines (see Baselines) using the “main database” and ONT sequencing data derived from bacterial isolates (see Data).

#### 2.3.1 Baselines

We selected five taxonomic classifiers that are representative of different approaches. As alignment-based taxonomic classifiers methods designed for long-read technologies, we selected MetaMaps [8] and MEGAN-LR [7] (alignments to reference nucleotides were created with minimap2 [24] as proposed in [25] and named as MEGAN-LR-nuc in that work). According to two existing taxonomic classification benchmark analyses using long reads, MetaMaps delivered the lowest number of false positive results for pseudo-mock communities [4] and MEGAN-LR for empirical mock communities [25]. Although a taxonomy classifier for metagenomic contigs, we also included MMseqs2 (nucleotide-based database) [26] as this tool was included in previous benchmarking for long-read data [25]. As a representative of k-mer matching methods, we selected Kraken2 [10], a widely used tool that can be applied to long-read data as well [27]. Finally, as a representative of deep-learning methods, we selected GeNet [11] as it provides a reasonable tradeoff between training and inference time.

#### 2.3.2 Experimental setup

MetageNN was trained with a dataset having 8*×* coverage of genomes from the “main database” to predict the genus of origin for a read. For validation, we generated a dataset containing 200 sequences for each of the 516 genomes. To accommodate the complexity of the model that needed to be trained, MetageNN model’s capacity was approximately doubled in terms of the number of neurons per layer (fig. 5, a, green text). There were approximately 17 million samples in the training dataset. MetageNN was trained using four GPUs and converged (early stopping) after three days of training. To reduce training time and need for computational resources only 1kbp sequences (see Sequence length analysis) were used, and predictions on ONT reads were obtained by segmenting into non-overlapping 1kbp chunks and using a majority voting strategy (tie broken by highest mean prediction probability). Then, predictions with a probability score greater or equal to 0.5 were considered as classified. MetageNN’s performance could therefore be improved further by direct training on reads with different read lengths (see Sequence length analysis).

GeNet was trained on the same training dataset as used by MetageNN. GeNet, in this case, also relies on a majority voting strategy and a probability threshold cutoff (0.5 as default). We extended GeNet to have a similar capacity (number of parameters) as MetageNN so that the difference between the two methods is primarily in how they encode long reads. GeNet training converged in approximately six days with four GPUs. We fine-tuned the hyperparameters of GeNet and MetageNN using the 1*×* dataset (see Coverage analysis).

We built all non-machine-learning taxonomic classifier indices using the genomes in the “main database” and these tools were run using its default settings.

#### 2.3.3 Evaluation metrics

We report classification performance in terms of sensitivity, precision and F1 scores. Specifically for each taxon, sensitivity describes the proportion of reads originating from it that is classified as such; precision indicates the proportion of correctly classified reads for that taxa, and F1 is the harmonic mean of precision and recall. With this, we report the average sensitivity, precision and F1 score across bacterial isolates per dataset tested. We also employed Wilcoxon’s Rank Sum Test to compare MetageNN’s performance relative to the baselines and report statistical significance for these comparisons.

#### 2.3.4 Results for all genomes

In this section, we provide overall results involving both test datasets (“Species in the database” and “Species out of the database”). This scenario represents what may occur in metagenomic sequencing experiments in which species with reference genomes and those without are both present, though the relative proportions may vary. fig. 3 (a) presents the results. For F1 and sensitivity scores, MMseqs2 achieved the highest average with 0.85 and 0.81, respectively, followed by Kraken2 with 0.83 and 0.79, respectively. MetageNN presented significantly higher F1 and sensitivity scores when compared to GeNet. For precision MEGAN-LR presented the highest scores, while MetageNN surpassed GeNet.

**Figure 3.**
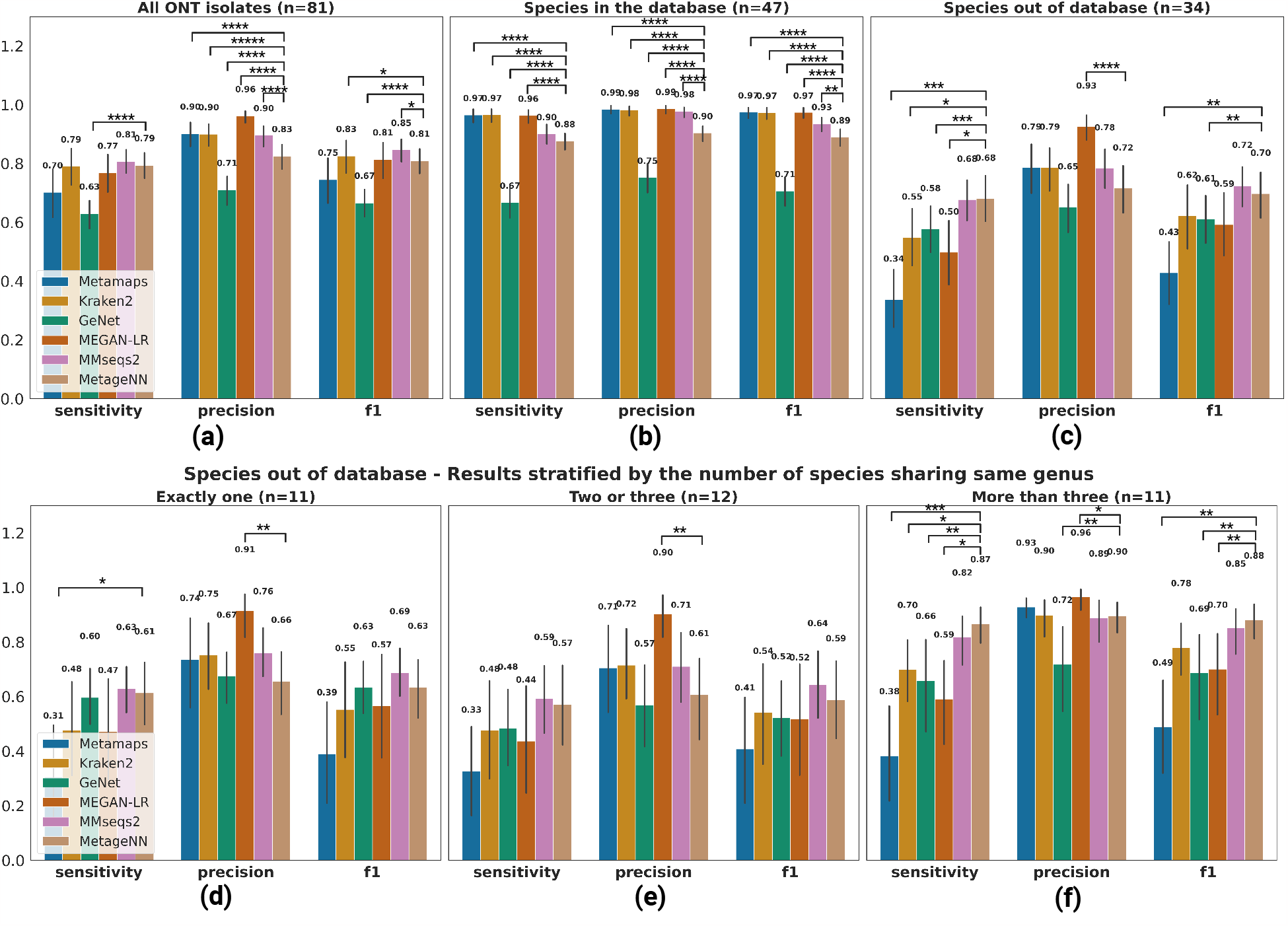
Results at the genus level of taxonomic classification methods applied to ONT data. Bar plots (error bars represent standard deviation across ONT bacterial isolates) showing results for MetaMaps, Kraken2, GeNet, MEGAN-LR, MMseqs2 and MetageNN. Average values of sensitivity, precision and F1 are shown on the top along with statistical significance bars on top, where * stands for digits after the decimal p-value point i.e., “****” signifies 1e-4. (a) Results aggregating all ONT isolates tested. (b) Results for the “Species in the database” dataset. (c) Results for the “Species out of the database” dataset. **Bottom: Results stratified by the number of species sharing the same genus**. Results for (d) “exactly one” (e) “two or three” and (f) “more than three” groups based on the “Species out of the database” dataset.

#### 2.3.5 Results for “Species in the database” settings

Here we present results based on the ONT read dataset “Species in the database” (described in Data). Results are presented in fig. 3, b.

As supported by previous benchmarking experiments [4, 28], alignment and k-mer-based classification methods are effective in the scenario where the correct reference genome of interest is included in the database. Kraken2, MetaMaps, and MEGAN-LR presented the best average F1, sensitivity, and precision scores, outperforming MetageNN, while MetageNN significantly improved over GeNet in all metrics tested.

#### 2.3.6 Results for “Species out of the database” settings

Frequently the correct genome of interest may not be part of the reference database, and a classifier’s ability to appropriately handle this scenario is an important attribute (i.e, a taxonomic classifier tool may be able to identify its rank above, in the case of a species, its genus). In contrast to existing conventional tools, we hypothesize that MetageNN offers an advantage in this regard through its use of the generalization properties of neural networks and its formulation of using short k-mer profiles of 6mers, which can help learn features shared across members of a taxonomic group. Therefore, we report the results (fig. 3 c) on the test dataset “Species out the database”.

Overall, MetageNN presented the highest sensitivity scores along with MMseqs2, significantly improving over MetaMaps, Kraken2, GeNet and MEGAN-LR by 100%, 23%, 17% and 36%, respectively. In addition to significantly improving over MetaMaps and GeNet in F1 scores, MetageNN presented higher F1 scores than most tools, except MMseqs2. The precision scores for MetageNN were slightly lower (non-significant results except for MEGAN-LR) than those for MetaMaps, Kraken2, and MMseqs2, but higher than those for GeNet.

To further characterize these results, we stratified them into three groups based on the number of references available for the taxa being tested (fig. 3, bottom). We noted that all tools had a lower performance for groups having three or fewer references (fig. 3 d and e). For the strictest setting of “exactly one” species sharing the same genus, MetageNN and MMseqs2 presented the highest F1 and sensitivity scores (with MetageNN significantly outperforming MetaMaps). Results for precision displayed MEGAN-LR outperforming all tools. For the group of “two or three”, the same trend of results was observed (fig. 3, e). Following that, all methods had their respective best results in the setting where there were “more than three” species per genus (fig. 3, f). Potentially due to its generalization ability and reliance on short k-mer profiles that might be shared across genera, MetageNN again achieved the best results, having the highest F1 and sensitivity scores of 0.88 and 0.87, respectively (significantly higher when compared to MetaMaps, GeNet and MEGAN-LR for both metrics and when compared to Kraken2 for sensitivity scores).

In summary, MetageNN has notable utility in settings where the correct genome might not be in the database, providing an alternative to conventional tools such as MetaMaps, Kraken2 and MEGAN-LR which have lower sensitivity in these cases.

### 2.4 MetageNN is more sensitive than MetaMaps, Kraken2, GeNet and MEGAN-LR at the community-level for ONT pseudo-mock communities

This section focuses on evaluating taxonomic classifiers on ONT pseudo-mock community samples. These communities, in contrast to the bacterial isolates in the previous section, are used to simulate metagenomic sequencing experiments and to report results at the community level (also known as detection-level) [4, 25].

#### 2.4.1 Experimental Setup

In order to obtain community-level results, we created ONT pseudo-mock communities from the reads of the bacterial isolates. Unlike experimental communities such as ZymoBIOMICS Microbial Community Standards ^1^ with only 10 species, using pseudo-mock datasets we can develop distinct community profiles with an increased and diverse number of species. We created 30 pseudo-communities consisting of ONT reads from species out of the database. The number of species in each pseudo-mock was randomly selected between 20 and 34 species in the “Species out of the database” list and we sampled a maximum of 100000 reads log-distributed across the species (to reduce computational resources for some tools). We refer to this dataset as “Mock - Species out of DB”.

We tested the same tools used in Baselines (except GeNet due to its poor performance presented in fig. 3) in the 30 pseudo-mock communities datasets. Following [25], we reported results using two approaches. The first one is the percentage of reads classified per tool (read classification). The second one is sensitivity, precision and F1 scores at the community level for the genus rank (i.e., the presence or absence of a genus based on a predefined threshold of cumulative read counts). We used a percentage threshold of 0.001, 0.1 and 1% of the total number of reads in each dataset (which provides approximately 1, 100, and 1000 minimum reads per threshold). We also employed Wilcoxon’s Rank Sum Test to compare MetageNN’s performance relative to the baselines and report statistical significance for these comparisons.

#### 2.4.2 Read classification results

We report the average percentage of reads classified for all mocks in fig. 4, a. Here, MetageNN outperformed all existing tools classifying 93.26% of the reads from species out of the database at the genus level. This result was followed by MMseqs2, Kraken2, MEGAN-LR and MetaMaps with 85.49%, 64.90%, 53.06% and 37.30%, respectively.

**Figure 4.**
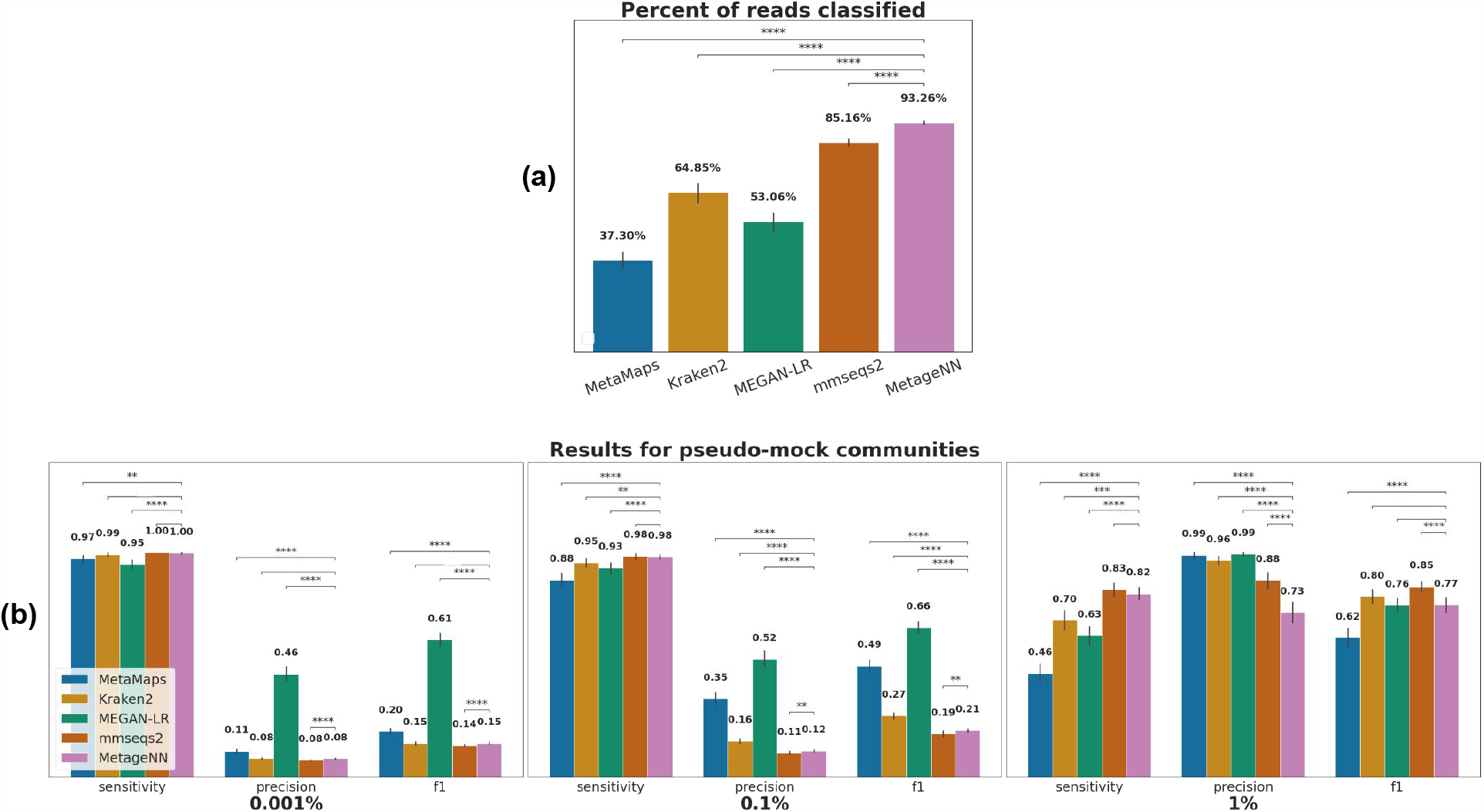
Results at the community-level (genus) of taxonomic classification methods applied to ONT pseudo-mock community data. (a) Average percentage of reads classified per tool for all mocks. (b) Bar plots showing results for MetaMaps, Kraken2, MEGAN-LR, MMseqs2 and MetageNN. Average values of sensitivity, precision and F1 are shown on the top. Statistical significance bars are displayed on top, where * stands for digits after the decimal p-value point i.e., “****” signifies 1e-4.

**Figure 5.**
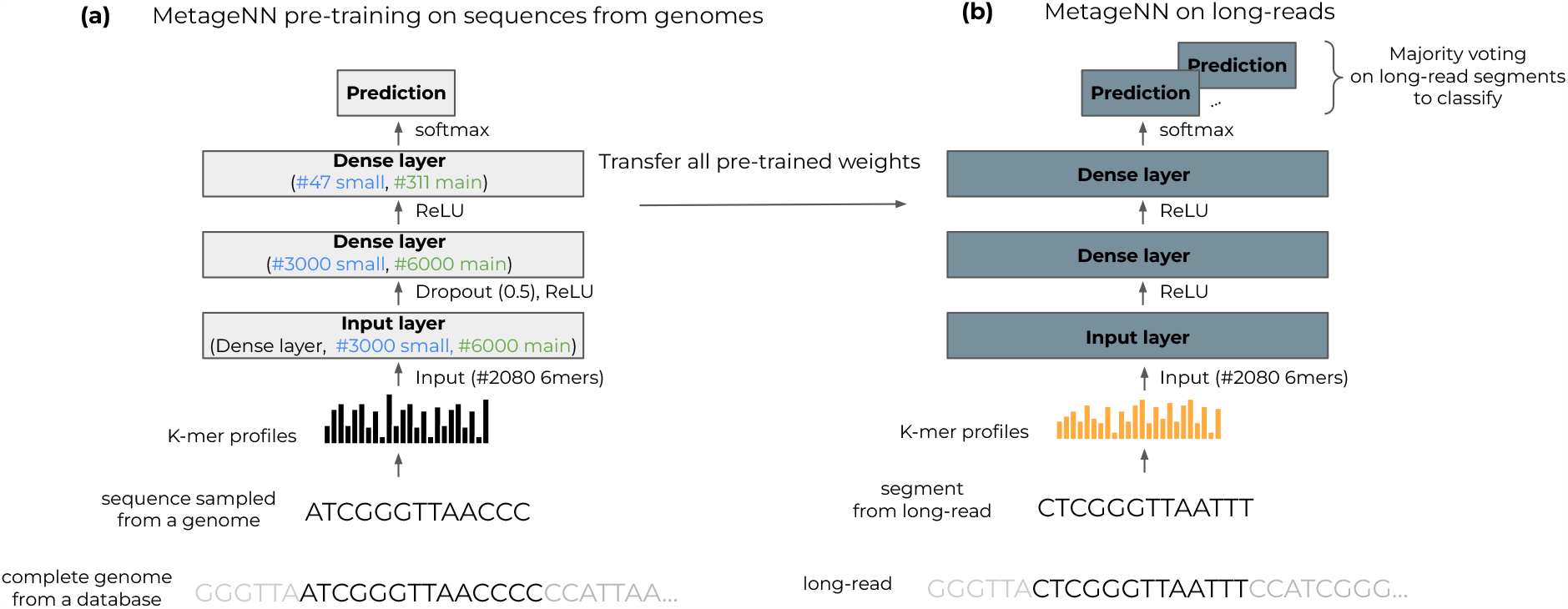
MetageNN framework for pre-training on long sequences from genomes and its direct application to long read data. (a) MetageNN utilizes the extensive collection of reference genomes available to sample long sequences. As a means of dealing with a different distribution of noisy long reads, MetageNN relies on computing short-k-mer profiles (6mers), which are more robust to sequencing errors and are used as input to the MetageNN architecture (gray rectangles represent layers). Parameters in blue are used for the small database and parameters in green for the main database (see Databases). A textual representation of activation functions and dropouts is provided between layers. (b) Once MetageNN training is completed, its learnt features are expected to be more robust to sequencing noise and thus can be directly transferred to long-read data (no fine-tuning needed). The classification of long-read data is based on a majority voting strategy.

#### 2.4.3 Community-level results

We report in (fig. 4, b) the average sensitivity, precision and F1 scores at the community-level for all pseudo-mock communities used. In this setting, MetageNN presented higher sensitivities (along with MMseqs2) for all thresholds and statistically outperformed MetaMaps and MEGAN-LR in all thresholds and Kraken2 for 0.1 and 1% thresholds. In terms of precision, MEGAN-LR delivered the best results at all thresholds, with MetageNN outperforming MMseqs2 at 0.001 and 0.1%. Finally, for F1, MEGAN-LR delivered the best results for 0.001 and 0.1% with MMseqs2 showing the optimum F1 score for the 1% threshold, while MetageNN surpassed MetaMaps. These results are in agreement with (fig. 3, c) and demonstrate the utility of MetageNN in situations where the exact genome reference is not available, since MetageNN can detect more reads in this case.

### 2.5 MetageNN is faster and requires less storage than alignment and k-mer-based tools, respectively

In this section, we analyzed the memory requirements to store its database and the classification speed obtained by all baselines. Results are depicted in table 1.

**Table 1:** Memory requirements for database storage and speed for each method on ONT data. Bold values indicate the best performer.

| Method | Memory (GB) | Speed (reads/sec) |
| --- | --- | --- |
| MetageNN | 0.839 | 1424 |
| GeNet | **0.758** | 134 |
| Kraken2 | 3.7 | **13471** |
| MetaMaps | 2.2 | 194 |
| MEGAN-LR | 5.1 | 606 |
| MMseqs2 | 27.8 | 588 |

To classify many organisms, conventional taxonomic classification methods require large databases and computational resources. When selecting a tool, the memory needed to store its database is a critical consideration. For example, Kraken2 requires a large amount of memory due to its formulation of storing pairs of the canonical minimizers of k-mers and their lowest common ancestors. As a result, Kraken2 showed a memory requirement to store its database of 3.7 GB. Similarly, MetaMaps uses a reference database of approximately 2.2 GB, followed by MEGAN-LR with 5.1 GB and MMseqs2 with 27.8 GB to store its indices. Since MetageNN is a neural network model, its final product is its learned weights. In this context, MetageNN needed 0.839 GB, decreasing the memory requirements for database storage by 77%, 61%, 83% and 96% compared to Kraken2 and MetaMaps, MEGAN-LR and MMseqs2 respectively. GeNet presented a similarly compact model with a memory requirement of 0.758 GB.

We also focussed on another important aspect of taxonomic classification tools i.e. classification speed. Due to the large volume of data produced by high-throughput sequencing technologies, this is an important feature to consider. In this study, MetageNN and existing tools were evaluated regarding the number of reads processed per second. We allowed four CPUs to be used by all tools as a representative of a commonly available system. Kraken2, including its database loading time, delivers the highest number of reads processed per second, with 13471 reads per second. MetageNN presented a rate of 1424 reads per second, including its k-mer counting and dataset loading time. MetaMaps, a mapping tool, delivers a speed of 194 reads per second, followed by MEGAN-LR and MMseqs2 with 606 and 588 reads per second. GeNet, a CNN-based model, processed 134 reads per second, providing the slowest tool among the ones tested, perhaps due to its execution on CPUs [29]. When considering the use of GPUs, deep-learning-based tools can offer a much higher speed but potentially at a higher cost for computing time.

## 3 Discussion and Conclusion

In this work, we explored the use of a machine learning approach for taxonomic classification of long-read data, and the need to address several challenges including the lack of training data for all species and the need to be robust to sequencing errors. MetageNN uses short k-mer-profiles (6mers) and relies on the availability of large databases of genomic sequences to learn features that can be transferred to the taxonomic classification of reads from long-read technologies.

We aimed to use a large training dataset of long-read-like sequences from genomes, and thus an architecture that would provide a feasible training time, reduced computational resources, and adequate classification speed is required. As a result, we demonstrated the effectiveness of MetageNN’s architecture in comparison with more sophisticated existing deep-learning-based taxonomic classification methods. Furthermore, a taxonomic classification model should be robust to sequencing errors originating from long-read technologies. Using synthetic long-reads with different sequencing error rates, MetageNN surpassed existing deep-learning-based tools demonstrating its robustness to sequencing errors (fig. 2).

We further trained MetageNN using a large dataset of approximately 17 million sequences and performed a comprehensive evaluation against representative taxonomic classification methods using real ONT reads. As a whole, MetageNN yielded an F1 score that significantly improved over GeNet when evaluating over all genomes (fig. 3, a).We also demonstrated the utility of MetageNN when the reference genome of the organism of interest is unavailable. MetageNN was more accurate (highest sensitivity scores along with MMseqs2) in identifying taxa of interest relative to existing methods in our dataset of bacterial isolates (fig. 3, c). MetageNN also demonstrated the same trend when evaluated on pseudo-mock communities having a higher sensitivity to all thresholds employed (fig. 4, b). Compared to MMseqs2, another tool with high sensitivity, MetageNN is memory-efficient, faster (table 1), classified more reads (fig. 4, a) and is less prone to generate false positives (fig. 4, b, MetageNN displayed higher precision than MMseqs2 in more thresholds).

There are several possible ways in which MetageNN’s performance could be improved. Firstly, training using a range of read lengths would likely improve MetageNN’s performance further in the “main database” setting and could bridge the gap observed relative to Kraken2 and MetaMaps when the genome of interest is in the database (fig. 3, b). Combined with a feature selection step based on the initial list of kmers, this could further improve MetageNN’s performance. Another possibility is to bridge the distribution shift from genomes to error-prone reads by supplementing the training data with sequences that contain synthetically introduced errors during training. A further extension of this would involve applying domain adaptation ideas based on adversarial training [30] with bacterial isolate sequencing data being used for training.

Prior work for using machine learning in taxonomic classification either relied on testing with simulated reads [15, 31, 14] or did not show advantages compared to conventional k-mer-matching and mapping-based approaches and real data [11, 13]. MetageNN is, to the best of our knowledge, the first machine learning-based method that shows improvements relative to conventional tools with real long-read data, when assessing performance for genomes that are not in the database.

Due to its neural network formulation, MetageNN has a smaller memory requirement than conventional tools. However, MetageNN in its current unoptimized form was slower than Kraken2. There are some potential directions for improving these results, including pruning [32] and quantization [33], which would increase speed and decrease memory usage. For this proof-of-concept work, MetageNN was trained for hundreds of species, but future directions include expanding it to predict thousands of species and exploring embedding-based classification to maintain memory efficiency [34]. In addition, assuming access to appropriate computational resources, GPUs could also contribute to an increase in MetaGeNN’s classification speed by a considerable amount, and could thus be an important future direction to investigate.

Generally, MetageNN offers a useful tradeoff between good F1 scores (fig. 3, a), particularly for taxa for which we don’t have genomes, smaller memory footprint relative to conventional tools, and faster prediction speed compared to mapping tools. These results demonstrate the feasibility of using machine-learning-based methods to further improve taxonomic classification accuracy and sensitivity in challenging real-world settings. Additionally, this work complements previous bench-marking [4, 25] for long-read taxonomic classifiers by exploring how existing methods perform for reads of species out of the database, a direction not explored previously. Apart from demonstrating MetageNN’s utility when conventional tools are unable to classify reads, the databases and carefully selected test datasets used in this study can be used for benchmarking to advance the application of machine-learning techniques for taxonomic classification.

## 4 Methods

### 4.1 MetageNN model

Machine learning has gained considerable popularity over the past few decades, with several break-through results observed in settings with a wide range of test domains with similar distributions of training data [35]. In the present case, to train a machine learning-based taxonomic classification method, we would ideally train on long-read sequencing data from all organisms of interest. For microbial genomes, this is not feasible, as only a very small fraction of species have corresponding sequencing data, e.g. SRA^2^ has ONT data for only 489 unique bacterial species. In comparison, the RefSeq [36] database has complete genomes for *>* 6,800 unique bacterial species (May 2022), potentially allowing for the detection of many more species when used for training. An even larger set of species have genomes if we consider draft genomes and metagenome-assembled genomes [37]. Therefore, a major assumption behind MetageNN is that we can explore large genomic databases such as RefSeq, and leverage the information learned by transferring it for long-read taxonomic classification.

To overcome the distribution shift between relatively error-free sequences from genomes and reads from long-read technologies, we transfer the features learned from k-mer profiles (also known as k-mer counts) of sequences sampled from RefSeq and adapt them to potentially erroneous long reads. This adaptation is aided by the use of small values for *k* (e.g. 6), as such k-mer-profiles are more robust to sequencing errors (a single base error affects fewer k-mers). Furthermore, we hypothesize that using k-mer profiles can be a useful encoding strategy for long-read sequencing data as they are read-length agnostic [6].

As training for a large number of genomes and corresponding sequence data can be computationally intensive, we investigated a straightforward and effective architecture for MetageNN, i.e., a three-layered feed-forward neural network with canonical k-mer profiles as input (accounting for different strands of sequencing reads [38]. We used the k-mer counting algorithm developed in [39] to enable rapid generation of k-mer profiles). For the network, while the first two layers of the network are used to extract features, the third layer was used to do multi-class taxonomic classification. To learn non-linear patterns, each layer in MetageNN is followed by a ReLU activation function [40]. To avoid overfitting, dropout [41] was applied to regularize the network. The model was trained using cross-entropy loss as follows:

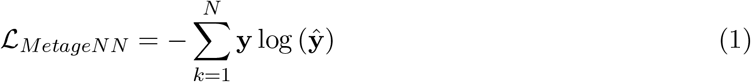

where **y** represents the true values and **ŷ** the predictions. fig. 5 schematically illustrates the workflow for MetageNN. Relative to methods with more complex network architectures, MetageNN offered significant improvements in inference speed, training time, and accuracy (see MetageNN is more effective and robust to sequencing errors than existing deep-learning-based taxonomic classification methods), and hence we used this simpler architecture for the proof-of-concept results using long-read sequencing data established in this study.

### 4.2 Data

#### 4.2.1 Test datasets

To evaluate MetageNN and existing tools on long-read sequencing data, we scanned NCBI’s Sequence Read Archive (SRA)^3^ for suitable ONT datasets (R.9.4.1 flow cell). For selecting bacterial isolates to download, we first identified those species with reference sequences (genomes) available in the NCBI RefSeq database^4^. We then confirmed that the strain name for the sequenced isolates had a corresponding match among the reference genomes available. Finally, we ensured that the match was correct by assembling (Flye [42] followed by two rounds of polishing with medaka^5^) the read data and verifying that the average nucleotide identity (ANI) to the reference was *>*99% (using the fastANI tool [43]). Following this process, we obtained ONT reads and corresponding genomes for 81 bacterial strains. In our settings, these strains belong to only one species in our databases used and for the remaining work, we refer to these strains as species or bacterial isolates. With this curated and verified list of ONT data, we developed two test datasets. The first dataset is to reflect the scenario in which the genome for a species is in the reference databases. We included reads from 47 species (out of 81) in this dataset that we refer to as the “Species in the database” dataset. The second dataset simulates the scenario in which the test reads are from a species whose genome is not in the reference database, referring to this as the “Species out of the database” dataset. In this case, we used the reads from the remaining 34 isolates. To reduce the computational resources required, we subsampled reads to have around 20000 per bacterial isolate used. When evaluating taxonomic classifiers, these bacterial isolate-derived test datasets offer a controlled environment for obtaining ground truth in different settings.

#### 4.2.2 Databases

To perform a fair evaluation, all baselines used the same database of genomes. Here, we devised two lists of genomes to include in their respective databases. We refer to the first database as the “small database” and it includes only genomes of the 47 species (47 genomes) listed in the “Species in the database” in the previous section (Test datasets). The purpose is to test various design choices for MetageNN’s model (see Setting parameters for the MetageNN model), to tune the hyperparameters for it, and to benchmark against existing deep-learning taxonomic classification methods which might otherwise be too computationally expensive to train and test on larger databases.

We refer to the second database as the “main database”. Here, we expanded the initial list of genomes from the “small database” approximately 10 times by randomly selecting additional genomes from the RefSeq representative genomes list. This resulted in a list of 516 genomes (belonging to 505 species and 311 genera). We devised this database to be used by all taxonomic classification methods to report the results when tested for “Species in the database” and “Species out of the database”.

## Authors’ contributions

R.P.S, C.S and N.N conceived the idea and the experiments. R.P.S designed and implemented MetageNN, conducted the evaluation and the experiments and wrote the manuscript. N.N and C.S reviewed the manuscript. All authors read and approved the final manuscript.

## Availability of data and materials

MetageNN’s model source code and the list of reference genomes and bacterial isolates data used are available at https://github.com/csb5/MetageNN.

## Footnotes

1 https://lomanlab.github.io/mockcommunity/

2 https://www.ncbi.nlm.nih.gov/sra

3 https://www.ncbi.nlm.nih.gov/sra

4 https://www.ncbi.nlm.nih.gov/refseq/

5 https://github.com/nanoporetech/medaka

## References

[1] Chiu, C. Y. & Miller, S. A. Clinical metagenomics. Nature Reviews Genetics 20, 341–355 (2019). URL 10.1038/s41576-019-0113-7.

[2] Wang, J. & Jia, H. Metagenome-wide association studies: fine-mining the microbiome. Nature Reviews Microbiology 14, 508–522 (2016). URL 10.1038/nrmicro.2016.83.

[3] Breitwieser, F. P., Lu, J. & Salzberg, S. L. A review of methods and databases for metagenomic classification and assembly. Briefings in bioinformatics 20, 1125–1136 (2019). URL https://pubmed.ncbi.nlm.nih.gov/29028872.29028872[pmid].

[4] Marić, J., Križanović, K., Riondet, S., Nagarajan, N. & Šikić, M. Benchmarking metage-nomic classification tools for long-read sequencing data. bioRxiv (2021). URL https://www.biorxiv.org/content/early/2021/08/18/2020.11.25.397729.

[5] Mande, S. S., Mohammed, M. H. & Ghosh, T. S. Classification of metagenomic sequences: methods and challenges. Briefings in Bioinformatics 13, 669–681 (2012). URL 10.1093/bib/bbs054. https://academic.oup.com/bib/article-pdf/13/6/669/482874/bbs054.pdf.

[6] Amarasinghe, S. L. et al. Opportunities and challenges in long-read sequencing data analysis. Genome Biology 21, 30 (2020).

[7] Huson, D. H. et al. Megan-lr: new algorithms allow accurate binning and easy interactive exploration of metagenomic long reads and contigs. Biology Direct 13, 6 (2018). URL 10.1186/s13062-018-0208-7.

[8] Dilthey, A. T., Jain, C., Koren, S. & Phillippy, A. M. Strain-level metagenomic assignment and compositional estimation for long reads with metamaps. Nature Communications 10, 3066 (2019). URL 10.1038/s41467-019-10934-2.

[9] Wang, Y., Zhao, Y., Bollas, A., Wang, Y. & Au, K. F. Nanopore sequencing technology, bioinformatics and applications. Nature Biotechnology 39, 1348–1365 (2021). URL 10.1038/s41587-021-01108-x.

[10] Wood, D. E., Lu, J. & Langmead, B. Improved metagenomic analysis with kraken 2. Genome Biology 20, 257 (2019).

[11] Rojas-Carulla, M. et al. Genet: Deep representations for metagenomics (2019).arXiv:1901. 11015.

[12] Liang, Q., Bible, P. W., Liu, Y., Zou, B. & Wei, L. DeepMicrobes: taxonomic classification for metagenomics with deep learning. NAR Genomics and Bioinformatics 2 (2020). URL 10.1093/nargab/lqaa009.Lqaa009.

[13] Vervier, K., Mahé, P., Tournoud, M., Veyrieras, J.-B. & Vert, J.-P. Large-scale machine learning for metagenomics sequence classification. Bioinformatics 32, 1023–1032 (2015). URL 10.1093/bioinformatics/btv683. https://academic.oup.com/bioinformatics/article-pdf/32/7/1023/19568386/btv683.pdf.

[14] Menegaux, R. & Vert, J.-P. Continuous embeddings of dna sequencing reads and application to metagenomics. Journal of Computational Biology 26, 509–518 (2019). URL 10.1089/cmb.2018.0174. PMID: 30785347, 10.1089/cmb.2018.0174.

[15] Georgiou, A., Fortuin, V., Mustafa, H. & Rätsch, G. Meta2: Memory-efficient taxonomic classification and abundance estimation for metagenomics with deep learning (2019). URL https://arxiv.org/abs/1909.13146.

[16] Wick, R. R. Badread: simulation of error-prone long reads. Journal of Open Source Software 4, 1316 (2019). URL 10.21105/joss.01316.

[17] Gregor, I., Dröge, J., Schirmer, M., Quince, C. & McHardy, A. C. PhyloPythiaS: a self-training method for the rapid reconstruction of low-ranking taxonomic bins from metagenomes. PeerJ 4, e1603 (2016). URL 10.7717/peerj.1603.

[18] Gehring, J., Auli, M., Grangier, D., Yarats, D. & Dauphin, Y. N. Convolutional sequence to sequence learning (2017). URL https://arxiv.org/abs/1705.03122.

[19] He, K., Zhang, X., Ren, S. & Sun, J. Deep residual learning for image recognition (2015). URL https://arxiv.org/abs/1512.03385.

[20] Amarasinghe, S. L. et al. Opportunities and challenges in long-read sequencing data analysis. Genome Biology 21, 30 (2020). URL 10.1186/s13059-020-1935-5.

[21] Rumelhart, D. E. & McClelland, J. L. Learning Internal Representations by Error Propagation, 318–362 (1987).

[22] Hochreiter, S. & Schmidhuber, J. Long short-term memory. Neural Comput. 9, 1735–1780 (1997). URL 10.1162/neco.1997.9.8.1735.

[23] Vaswani, A. et al. Attention is all you need. In Proceedings of the 31st International Conference on Neural Information Processing Systems, NIPS’17, 6000–6010 (Curran Associates Inc., Red Hook, NY, USA, 2017).

[24] Li, H. Minimap2: pairwise alignment for nucleotide sequences. Bioinformatics 34, 3094–3100 (2018). URL 10.1093/bioinformatics/bty191. https://academic.oup.com/bioinformatics/article-pdf/34/18/3094/48919122/bioinformatics_34_18_3094.pdf.

[25] Portik, D. M., Brown, C. T. & Pierce-Ward, N. T. Evaluation of taxonomic classification and profiling methods for long-read shotgun metagenomic sequencing datasets. BMC Bioinformatics 23, 541 (2022). URL 10.1186/s12859-022-05103-0.

[26] Mirdita, M., Steinegger, M., Breitwieser, F., Söding, J. & Levy Karin, E. Fast and sensitive taxonomic assignment to metagenomic contigs. Bioinformatics 37, 3029–3031 (2021). URL 10.1093/bioinformatics/btab184. https://academic.oup.com/bioinformatics/article-pdf/37/18/3029/40471479/btab184_supplementary_data.pdf.

[27] Leidenfrost, R. M., Pöther, D.-C., Jäckel, U. & Wünschiers, R. Benchmarking the minion: Evaluating long reads for microbial profiling. Scientific Reports 10, 5125 (2020). URL 10.1038/s41598-020-61989-x.

[28] Ye, S. H., Siddle, K. J., Park, D. J. & Sabeti, P. C. Benchmarking metagenomics tools for taxonomic classification. Cell 178, 779–794 (2019).

[29] Liu, Y. et al. Optimizing cnn model inference on cpus (2018). URL https://arxiv.org/abs/1809.02697.

[30] Ganin, Y. & Lempitsky, V. Unsupervised domain adaptation by backpropagation. In Bach, F. & Blei, D. (eds.) Proceedings of the 32nd International Conference on Machine Learning, vol. 37 of Proceedings of Machine Learning Research, 1180–1189 (PMLR, Lille, France, 2015). URL https://proceedings.mlr.press/v37/ganin15.html.

[31] Menegaux, R. & Vert, J.-P. Embedding the de bruijn graph, and applications to metagenomics. bioRxiv (2020).

[32] Blalock, D., Ortiz, J. J. G., Frankle, J. & Guttag, J. What is the state of neural network pruning? (2020). URL https://arxiv.org/abs/2003.03033.

[33] Gholami, A. et al. A survey of quantization methods for efficient neural network inference (2021). URL https://arxiv.org/abs/2103.13630.

[34] Wei, T., Mao, Z., Shi, J.-X., Li, Y.-F. & Zhang, M.-L. A survey on extreme multi-label learning (2022). URL https://arxiv.org/abs/2210.03968.

[35] Pan, S. J. & Yang, Q. A survey on transfer learning. IEEE Transactions on Knowledge and Data Engineering 22, 1345–1359 (2010).

[36] O’Leary, N. A. & Wright, M. W. Reference sequence (RefSeq) database at NCBI: current status, taxonomic expansion, and functional annotation. Nucleic Acids Research 44, D733–D745 (2015). URL 10.1093/nar/gkv1189. https://academic.oup.com/nar/article-pdf/44/D1/D733/9482930/gkv1189.pdf.

[37] Almeida, A. et al. A new genomic blueprint of the human gut microbiota. Nature 568, 499–504 (2019). URL 10.1038/s41586-019-0965-1.

[38] Bussi, Y., Kapon, R. & Reich, Z. Large-scale k-mer-based analysis of the informational properties of genomes, comparative genomics and taxonomy. PLOS ONE 16, 1–27 (2021). URL 10.1371/journal.pone.0258693.

[39] Gregor, I., Dröge, J., Schirmer, M., Quince, C. & McHardy, A. C. PhyloPythiaS+: a self-training method for the rapid reconstruction of low-ranking taxonomic bins from metagenomes. PeerJ 4, e1603 (2016).

[40] Nair, V. & Hinton, G. E. Rectified linear units improve restricted boltzmann machines. In Proceedings of the 27th International Conference on International Conference on Machine Learning, ICML’10, 807–814 (Omnipress, Madison, WI, USA, 2010).

[41] Srivastava, N., Hinton, G., Krizhevsky, A., Sutskever, I. & Salakhutdinov, R. Dropout: A simple way to prevent neural networks from overfitting. Journal of Machine Learning Research 15, 1929–1958 (2014). URL http://jmlr.org/papers/v15/srivastava14a.html.

[42] Kolmogorov, M. et al. metaflye: scalable long-read metagenome assembly using repeat graphs. Nature Methods 17, 1103–1110 (2020). URL 10.1038/s41592-020-00971-x.

[43] Jain, C., Rodriguez-R, L. M., Phillippy, A. M., Konstantinidis, K. T. & Aluru, S. High throughput ani analysis of 90k prokaryotic genomes reveals clear species boundaries. Nature Communications 9, 5114 (2018). URL 10.1038/s41467-018-07641-9.

